# A Novel, Versatile Speculum-free Callascope for Clinical Examination and Self-Visualization of the Cervix

**DOI:** 10.1101/618348

**Authors:** Mercy N. Asiedu, Júlia S. Agudogo, Mary Elizabeth Dotson, Marlee S. Krieger, John W. Schmitt, Megan Huchko, Gita Suneja, Rae Jean Proeschold-Bell, Jennifer S. Smith, Deborah Jenson, Wesley Hogan, Nirmala Ramanujam

**Affiliations:** Department of Biomedical Engineering, Duke University, Durham, North Carolina, United States of America; Center for Global Women’s Health Technologies, Duke University, Durham, North Carolina, United States of America; Duke Global Health Institute, Duke University, Durham, North Carolina, United States of America; Department of Obstetrics and Gynecology, Duke University, Durham, North Carolina, United States of America; Department of Radiation Oncology, Duke University, Durham, North Carolina, United States of America; Department of Epidemiology, University of North Carolina, Chapel Hill, North Carolina; Department of Romance Studies, Duke University, Durham, North Carolina, United States of America; Center for Documentary Studies, Duke University, Durham, North Carolina, United States of America

## Abstract

**Background:** Invasive cervical cancer is preventable, yet affects 500,000 women worldwide each year, and over half these women die. Barriers to cervical cancer screening include lack of awareness of cervical cancer and the cervix, fear of the speculum, and lack of women-centric technologies. We developed a low-cost (∼$50), cervix-imaging device called the Callascope, which comprises an imaging component, camera and inserter that eliminates the need for a speculum and enables self-insertion. We sought to assess the quality of physicians’ images of the cervix using the Callascope versus the speculum in live patients and study women’s willingness to independently use the Callascope to image their cervix.

**Methods:** We conducted two main studies: (1) a clinical study in which a physician imaged the cervix of patients using both the speculum and Callascope in a 2×2 crossover design; and (2) home-based self-cervix imaging with the Callascope.

**Results:** Participants of the clinical study (n=28) and home study (n=12) all indicated greater comfort and an overall preference for the Callascope over the speculum. The clinical study data indicated that the Callascope enabled similar visualization compared to the speculum while significantly improving patient experience. With physician insertion and manipulation, the Callascope enabled cervix visualization for 82% of participants. In the home-study, 83% of participants were able to visualize their cervix with the Callascope on the first try and 100% after multiple attempts.

**Conclusion:** The Callascope is more comfortable and provides similar visualization to the speculum. The Callascope can be used by medical providers for clinical exams while also enabling home self-screening for cervical cancer and promoting a better understanding of one’s cervix to increase awareness of cervical screening needs. The Callascope may increase cervical cancer screening rates through reducing barriers including cost, discomfort, lack of awareness and stigma.

## Introduction

In the U.S., the cervical care cascade begins with screening for cervical cancer using the Papanicolaou (Pap) smear combined with human papillomavirus (HPV) testing. Colposcopy, which visualizes the acetic acid-stained cervix with a low-power microscope, followed by biopsy of cervical abnormalities, serves as a confirmatory test for women with positive screening results. ^1^ Women with pre-cancer are treated via excisional or ablative treatments. Women with cancer are referred to a combination of local and/or systemic therapy, depending on the stage of invasive disease. This model is not practically implementable in medically underserved regions due to lack of resources to procure, implement, and maintain the technologies in the care cascade. Thus, alternative protocols that employ low-cost, simple-to-use technologies are needed to mitigate cervical cancer. ^2^ The most common screening tool in lower-resource countries is visual inspection with acetic acid (VIA) using the naked eye or a digital camera followed by either colposcopy (common practice in middle income countries) or treatment (most common in low income countries).^3^

Regardless of the setting, the duckbill speculum is used to visualize the cervix whether it is for a Pap smear, VIA or colposcopy. This is a significant factor in women avoiding cervical cancer screening, largely due to anxiety, fear, discomfort, pain, embarrassment, and/or vulnerability during the procedure. ^4-7^ In the U.S., Middle-aged African women have the highest incidence and mortality for cervical cancer. Their perceived pain of the procedure in combination with the high costs of office visits is associated with a six-fold increase in non-adherence to screening relative to the general population.^4^ In a longitudinal study of sexually active young women, poor compliance with return screening for Pap smears is again associated with perceptions of pain.^8^ These barriers to screening are echoed globally. In Australia, a study seeking to determine women’s attitudes towards physician versus self–insertion of the standard speculum found that 91% of 133 women would choose self-insertion over physician insertion, and that women felt discomfort, embarrassment and vulnerability from having another person insert a device and examine their cervix. ^9^ A study of 354 women in Moshi, Tanzania revealed that key behavioral barriers for cervical cancer screening were concerns about embarrassment and pain due to the speculum, and physician gender. ^10^ The speculum is a cause of discomfort, especially for women with vaginismus, a condition involving the involuntary tightening of the vagina often caused by sexual abuse. ^11^ It is indeed worth considering that the countries that typically have the highest sexual violence rates worldwide ^12,13^ also have the highest rate of cervical cancer incidence and mortality. ^14^ Rendering the screening process speculum-free when women initially enter the care cascade would allow the majority of women who do not have abnormalities to be screened without a speculum and would provide an opportunity to educate women who require referral about the need for speculum-based treatment once they are in the health system.

Other barriers cited are the medical and travel costs, and time associated with the need to visit a tertiary health institutions for cervical cancer screening.^15,16^ Even though there are efforts to deploy visual inspection to community health settings, most women still need a follow-up visit for confirmatory colposcopy and another follow-up visit for treatment. In this era of patient-centered medicine, self-imaging of the cervix would allow colposcopy to reach marginalized women and reduce the need for multiple hospital visits.^17^ Self-sampling of the cervix has previously been limited by the need for the speculum, to visualize the cervix and the cervical os for accurate sampling. HPV self-sampling overcomes these limitations through sensitive molecular testing and has proven feasibility of success.^17^ However, HPV sampling is not readily available and when it is, there is still a requirement for follow-up cervix visualization to improve specificity. Image capture with a low-cost, easy-to-use, self-imaging device, which can enable diagnostic reading by a remote health provider would allow women to visualize their cervices and receive expert feedback, without the need for a hospital visit. This could also be coupled with HPV self-sampling to more effectively triage women.

A third barrier is that in many settings, women are often not aware of their reproductive anatomy and health, which can deter positive health-seeking behavior, particularly as it relates to cervical cancer screening. ^18^ In 2014, the American Congress of Obstetricians and Gynecologists (ACOG) and Women’s Health magazine conducted a survey of 7,500 women and found that 64% of them could not identify a picture of the cervix. The same survey found that 54% of women admitted keeping secrets related to their reproductive anatomy from their gynecologists, perhaps out of embarrassment related to issues pertaining to their reproductive system. ^19^ Women’s awareness of their reproductive anatomy has been associated with high education level of both the women and their parents, exposure to media outlets and living near healthcare facilities.^20-22^ Further, several studies have found that educating women on their cervix and cervical cancer significantly increased cervical cancer screening rates more than two-fold. ^18,19,23-29^ Simply put, giving women access to accurate information and a comprehensive education on their sexual anatomy can significantly improve their understanding of sexual and reproductive health. One strategy to empower women to care and advocate for their reproductive autonomy is the provision of self-exploration technology that facilitates women’s visualization, understanding and appreciation of their own vagina and cervix.

We have previously published work on a low-cost, speculum-free, cervical visualization device, the Callascope. ^30^ We have also developed an android-based mobile application HIPPA (Health Insurance Portability and Accountability Act of 1996)-compliant image capture and cloud storage. The Callascope consists of an inserter and a 2-megapixel (MP) off-the-shelf camera (Supereyes Y002) that fits in a channel in the inserter to enable image capture. Standard-of-care speculums have bills that are opened once inserted into the vagina to expand the entire vaginal canal and a slightly longer lower lip used to manipulate the cervix (**Fig. 1a,b**) to center the os, the canal that leads to the uterus. The Callascope inserter, referred to as the Calla, has a slim tubular body the size of a small tampon, and a funnel-like curved tip, shaped like a Calla Lily that is used to part only the vaginal walls closest to the cervix, with a lip for cervix manipulation to center the os (**Fig. 1c,b**). Centering the os is important since cervical pre-cancers originate and spread from this area of the cervix. It is therefore important to obtain an image where the os and the area around it can be seen. Mechanical finite element testing simulations, testing on custom vaginal phantoms, and evaluation on healthy volunteers demonstrated that this technology can serve as an effective alternative to the speculum for visualization of cervices with different orientations (due anteverted, retroverted, and sideverted uteri). ^30^ In addition, questionnaire responses from volunteers indicated that > 92% of them preferred the Calla compared to the traditional speculum. 30

**Figure 1.**
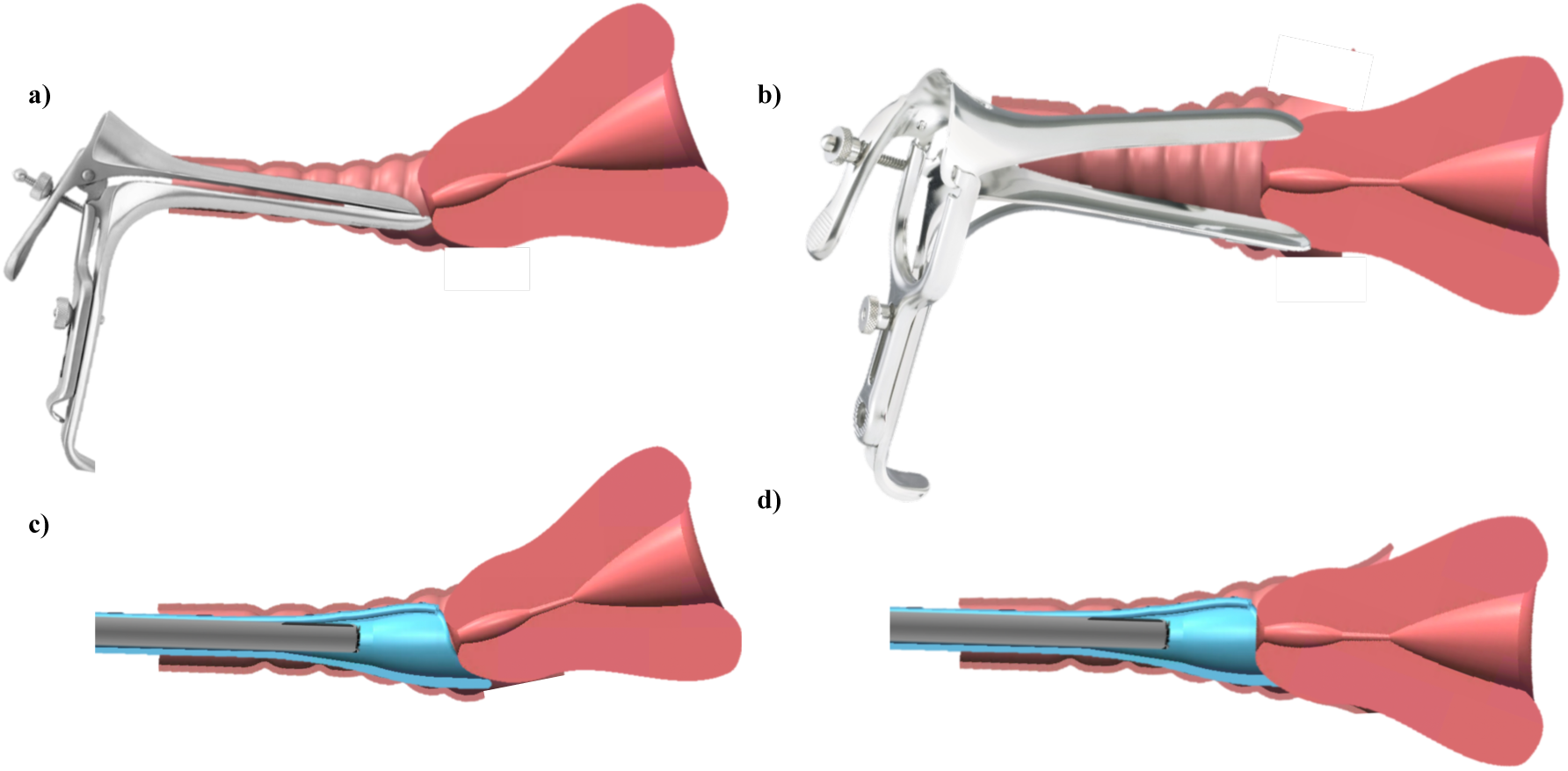
a) Standard-of-care speculum inserted into vagina with tilted uterus, b) Speculum bill centering the cervix for imaging, c) Callascope inserted into the vagina with tilted uterus, d) Callascope lip centering the cervix for imaging.

This manuscript builds upon our previous work and reports on three distinct avenues of inquiry that directly serve to address the barriers described in the introduction: (1) a pilot clinical study to demonstrate the effectiveness of the Callascope for examination of the cervices of patients undergoing cervical cancer screening, (2) healthy volunteer studies where women performed a home-based self-exam after completion of a short training session in a clinical setting to demonstrate the feasibility of self-imaging in the absence of a health professional, and finally (3) pre-and post-surveys as well as audio reflections (in the self-imaging group only) to collect perceptions of the speculum, cervical cancer screening, and experiences with the Callascope. The self-exam investigation differed from our previous healthy volunteer study in that women imaged their cervices independently, without a physician or health provider present and further used the device several times in their homes over a period of one week. The clinical study extends the capabilities of the Callascope over what was published previously and used a 2×2 crossover design to directly compare cervical visualization with the Callascope versus the speculum on the same patients. Visualization with the Callascope with physician use and self-imaging produced comparable results-the Callascope visualized the cervix in more than 80% of women in both groups. Further, this increased to 100% in the self-imaging group when they were allowed to repeat this at least once at home. Both patients and healthy volunteers indicated overwhelming preference for the Callascope over the standard-of-care speculum. In their audio reflections from the home study, volunteers indicated that the Callascope was easy to use and they expressed improved awareness of their reproductive anatomy. Overall these studies suggest the feasibility of using the Callascope for speculum-free clinic-based imaging and self-imaging of the cervix.

## Methods

### Studies overview

This paper presents two sub-studies, each with a distinct group of participants: (1) a physician-administered exam with the Callascope on patients undergoing a routine Pap smear; and (2) self-imaging of the cervix with the Callascope over a period of one week at home after initial training in a clinical setting. This study was approved at the Duke University Medical Center Institutional Review Board (DUMC IRB) (Pro00008173). All participants provided written informed consent prior to participating in these studies. High-level disinfection of the Calla prior to clinical and self-exams was performed by submersion in 2% hydrogen peroxide for 8 minutes at room temperature, as recommended for semi-critical devices (contact with mucous membranes or non-intact skin during patient use) ^31,32^ while the imaging component was wiped down with a Sani wipe. Components that came into direct contact with vaginal mucosa were discarded after use and were not shared between participants. A summary of data collected from surveys in both studies is found in **Table 1**.

**Table 1:** Summary of data collected from surveys in two distinct studies.

| **Table 1: Summary of data collected from surveys in two distinct studies** |  |  |
| --- | --- | --- |
| **Study** | **Surveys/interviews** | **Data collected** |
| Study 1: Clinical cervix imaging | Pre-insertion surveys | Demographics, top features for screening, perceptions of the speculum and Callascope |
|  | Post-insertion surveys | Discomfort comparison between speculum and Callascope on the Wong Baker Pain Scale. |
| Study 2: Self-imaging of the cervix | Pre-insertion surveys | Demographics, top features for screening, perceptions of the speculum and Callascope |
|  | Post-exam First Insertion survey (in clinic) | Ease of use and discomfort on a Likert Scale |
|  | Post-exam Last insertion survey (at home) | Ease of use and discomfort on a Likert Scale |
|  | Audio reflections | In-depth thoughts on the Callascope self-exam |

### Study 1: Clinical exam

The clinical study was performed to determine the comfort and visualization capabilities of the Callascope in comparison to the duckbill speculum in a 2×2 cross over study. The 2×2 cross over study was chosen to reduce patient bias from discomfort associated with order of placement. It also served to reduce physician bias from prior knowledge of the cervix position. Adult patients (n=28) undergoing routine Pap smears were recruited for the study. Pap smear patients were chosen due to the logistical challenges of performing a 2×2 crossover study during colposcopy in which multiple procedures (contrast application, biopsy, endocervical curettage, etc.) are being carried out. Patients were asked to complete a pre-exam survey. Per the crossover design, either the speculum or the Callascope was randomly assigned in terms of order of use; n=17 of the participants were examined with the speculum first followed by the Callascope, and n=11 participants were examined with the Callascope first, followed by the speculum. There was a two-minute delay between insertions to reduce any carryover effect. It should be noted in order to maintain consistency in imaging, the Callascope was used either alone (speculum-free imaging) or through the duckbill speculum (speculum-based imaging). The Pap smear was performed after imaging was completed with both approaches. After the procedure, the participant was asked to fill out a post-exam survey which assessed comfort, quantified on a 0-10 Wong-Baker pain scale, with the Callascope and the speculum.

### Study 2: Self-exam

The self-exam study involved training healthy volunteers to use the Callascope and usage of the device over a one-week period to assess ease-of-use and feasibility of imaging the cervix without physician guidance. This involved (1) on-site training and initial self-imaging of the cervix and (2) repeated self-imaging at home.

### Part 1: On-site training and initial cervix imaging

Healthy volunteers (n=12), were recruited using online advertisements and flyers, and scheduled for a research visit at DUMC. Following informed consent, participants were asked to complete a pre-insertion survey (**Supplemental fig. 2**). Participants were then given a user kit containing a Callascope, an android phone and phone charger, Sani wipes, vaginal wipes, lubricating jelly, printed user guide, and audio reflection guide (**Supplementary fig. 3**). Participants were asked to watch a <u>video tutorial</u>, available on the custom Calla mobile application, which provided information on how to assemble the Callascope and use it to capture images of the cervix (**Supplemental Fig. 4**). Training took 5-10 minutes. After watching the tutorial, participants could ask the study coordinators questions regarding use of the Callascope. Participants were then allowed to capture an image of their cervix with the Callascope in a private room. This typically took 5-10 minutes. A study nurse confirmed whether or not the images captured were those of the cervix. After the self-exam, participants were asked to indicate their level of discomfort on a Likert scale with the following options: “No discomfort”, “Slight discomfort”, “Moderate discomfort”, “A lot of discomfort”, and “Extreme discomfort”. They also indicated how easy or difficult it was to follow the instructions and to use the Callascope to find and manipulate their cervix.

### Part 2: Home self-exam

After completing their on-site training and initial self-imaging session, the participants were sent home with the Calla user kit for a week. At home, participants were asked to perform a self-visualization exam at least once and up to 3 times and to capture images of their cervix using the custom mobile application which enables them to rewatch the video tutorial, capture images, and store images. At the end of the one-week period, participants completed a final survey to assess the degree of ease/difficulty of use and comfort/discomfort level on a Likert scale, after using the Callascope during their final attempt at home. Participants also completed an audio reflection guide to share their thoughts on exploring their inner reproductive parts for the first time and to give feedback on the study. In the event that participants menstruated during the study, they were asked to postpone imaging until the end of their period.

### Statistical analysis

A two-tailed Student’s t-test was used to determine if any significant differences were observed in visual area when the cervix was viewed with the speculum versus the Callascope, with the null hypothesis that there was no significant difference in visual area (alpha=0.05). A two-tailed Student’s t-test was also used to analyze post-exam survey data comparing comfort/discomfort between the speculum and Callascope, with the null hypothesis that the Callascope enabled the same comfort level as the speculum (alpha=0.05). A paired, one-tailed t-test (alpha=0.05) was used to determine significant differences in post-exam responses for comfort/discomfort and ease-of-use between the initial Callascope use (during the training exam) and the final use (during the home exam).

## Results

Pre-insertion survey responses by substudy are summarized in **Table 2**. Participants represented different races and had a wide range of ages, vaginal births and BMIs. Seventy five percent of women in the self-exam group regularly used tampons/menstrual cups whereas only 40% of women from the clinical study regularly used tampons or menstrual cups. Most women who participated in the clinical study reported having had greater than ten speculum exams, while all participants in the self-exam group had at least one previous speculum exam. Results from the pre-insertion survey found that 50% of women from the self-exam group and 62.5% of women from the clinical study found the speculum to be a barrier to cervical cancer screening. Based on appearance only, more women from the self-exam and clinical study groups were willing to use the Callascope over the speculum. Participants in the self-exam group ranked comfort as third most important (cost and adequate assessment of cancer risk ranked higher), while those in the clinical study ranked comfort as most important. Procedure and travel time were least important for participants in the self-exam group while physician gender was least important to the clinical study group.

**Table 2:**
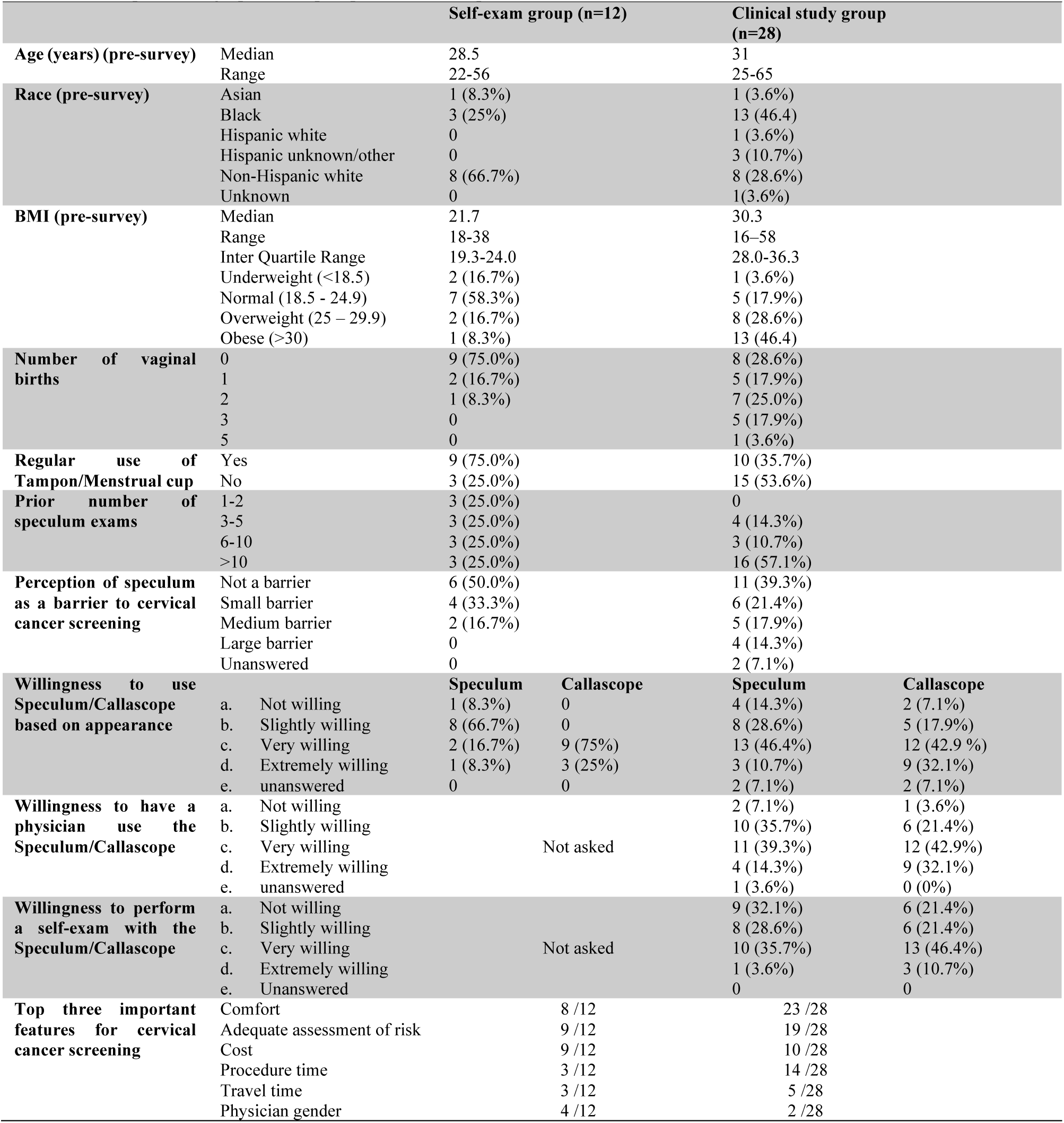
Participant demographics and pre-questionnaire responses.

### The Callascope has equivalent visualization but significantly superior patient comfort relative to the speculum

The Callascope was evaluated in a clinical study on 28 patients undergoing a Pap smear at DUMC to evaluate cervical visualization and patient acceptability. Patient demographics for the study are outlined in **Table 2**. Prior to the exam, patients filled out pre-questionnaires on the Callascope and the speculum. Participants in the clinical study ranked comfort, adequate assessment of risk and cost as the top 3 most important features for cervical cancer screening (**Table 2**). More than half (62%) of patients perceived the speculum as a barrier to screening and a higher percentage indicated either very or extreme willingness to use the Callascope for both self-(57.1%) and clinical-(75%) exams over the speculum (39.3% for self, and 53.6% for clinical) (**Table 2**). Representative images from the speculum and Callascope are shown in **Fig. 4(a)**. Visual area calculated for the speculum and Callascope inserter is shown in **Fig. 4(b)** with no significant differences between the speculum and the Callascope for lower BMIs, but a trend towards slighter lower visualization area with the Callascope in women with BMI > 25. With physician insertion, the Callascope enabled visualization of the cervix in 23/28 (82.1%) of women. The women for whom the Callascope failed, were either overweight (BMI>25) or obese (BMI >30). In these instances, a larger speculum size was used, whereas the same size of Callascope was used. We can easily overcome this limitation by introducing two different sizes of the Callascope. In the post-insertion questionnaire, the Callascope was scored by patients as causing significantly less discomfort than the speculum for insertion, manipulation, and removal (**Fig. 4(c**)). These results demonstrate that the Callascope can be readily used for clinical cervix and vaginal wall visualization in place of the standard-of-care speculum, while enabling significantly improved patient experience.

**Figure 4.**
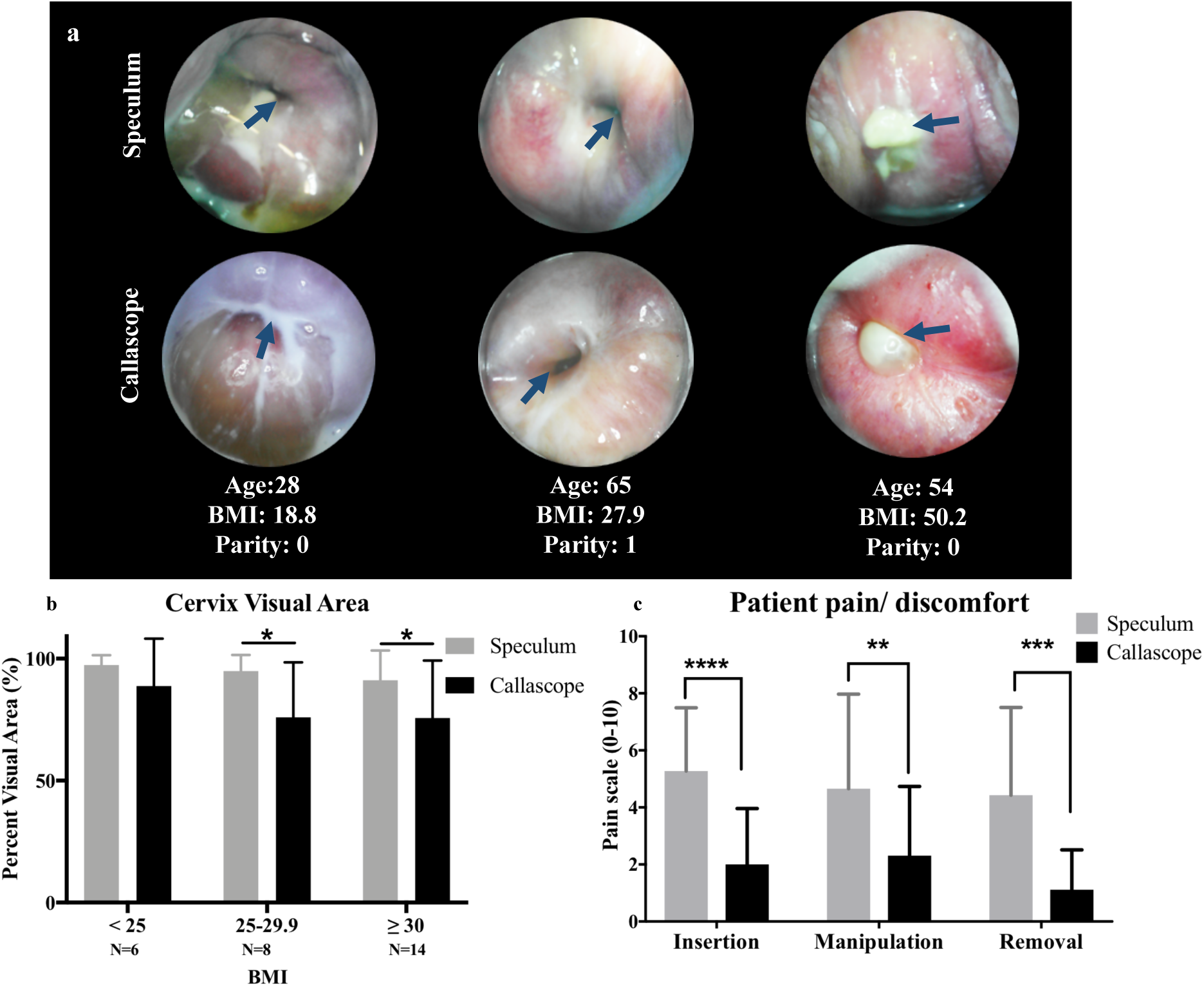
Cervix imaging results from clinical feasibility studies comparing Callascope to the standard-of-care speculum a) Representative images from clinical feasibility studies comparing Callascope to the speculum with age, BMI, and parity indicated. Images from both the Calla and Speculum are cropped to only show the cervix without vaginal walls, or device parts. The speculum and Callascope images from the same patient are shown for each participant. Blue arrows point to the cervical os. Blood in images are primarily from Pap smear sample removal or menstruation. b) Cervix visual area results comparing Callascope to speculum by BMI. It should be noted that different speculum sizes were used depending on patient BMI and parity, i.e., a larger Graves speculum was used on women with BMI > 24.9 whereas the same Callascope size was used across all BMI ranges. c) Patient discomfort for Callascope compared to speculum. P-values are *=0.05, **=0.005, ***=0.0005, ****=0.00005. Error bars are standard deviation

### Self-imaging of the cervix is comparable to physician-based visualization with the Callascope

In the self-examination study, the study nurse confirmed that 10 out of 12 (83.3%) participants were able to visualize their cervix on their first attempt. Of the 12 participants, 11 (91.7%) were also able to use the device to visualize their cervix at home; one participant was unable to do so due to a self-reported device fail with the camera of the Callascope. All participants were able to capture at least one image of the cervix by the end of the study. Images of the cervix captured in the self-exam in the clinic and at home are shown in **Fig. 4a**. Additionally, a representative video of the insertion of the Callascope, showing the vaginal wall and cervix manipulation, is shown in <u>this link</u>.

The post-insertion survey results from the self-exam in the clinic (initial) and self-exam at home (final) are shown in **Fig. 5e-l**. To summarize, a paired, two-tailed t-test (alpha=0.025) demonstrated no significant differences in discomfort or ease-of-use between the initial and final Callascope use, although the sample size was small. Most of the participants found the instructions easy to use and half of the participants found it extremely easy to slightly easy to find their cervix and this improved with repeat insertions at home (**Fig. 5e,f**). Most women found it easy to visualize their cervix with the Callascope but the difficulty they reported did increase when the exam was performed at home vs. the clinic (**Fig. 5 g,h**). Almost half (45%) of women had little to no discomfort inserting the Callascope during the initial use, 46% found insertion moderately uncomfortable and only 9% (1 participant) had a lot of discomfort during insertion (**Fig. 5i**). In the case of final use, the same participant continued to report a lot of insertion discomfort, but 67% reported little to no discomfort (**Fig. 5j**). All participants found that removal of the Callascope posed no discomfort or only slight discomfort (**Fig. 5k,l)**. In the final post-insertion survey, all of the participants indicated that they would recommend the Callascope to others.

**Figure 5.**
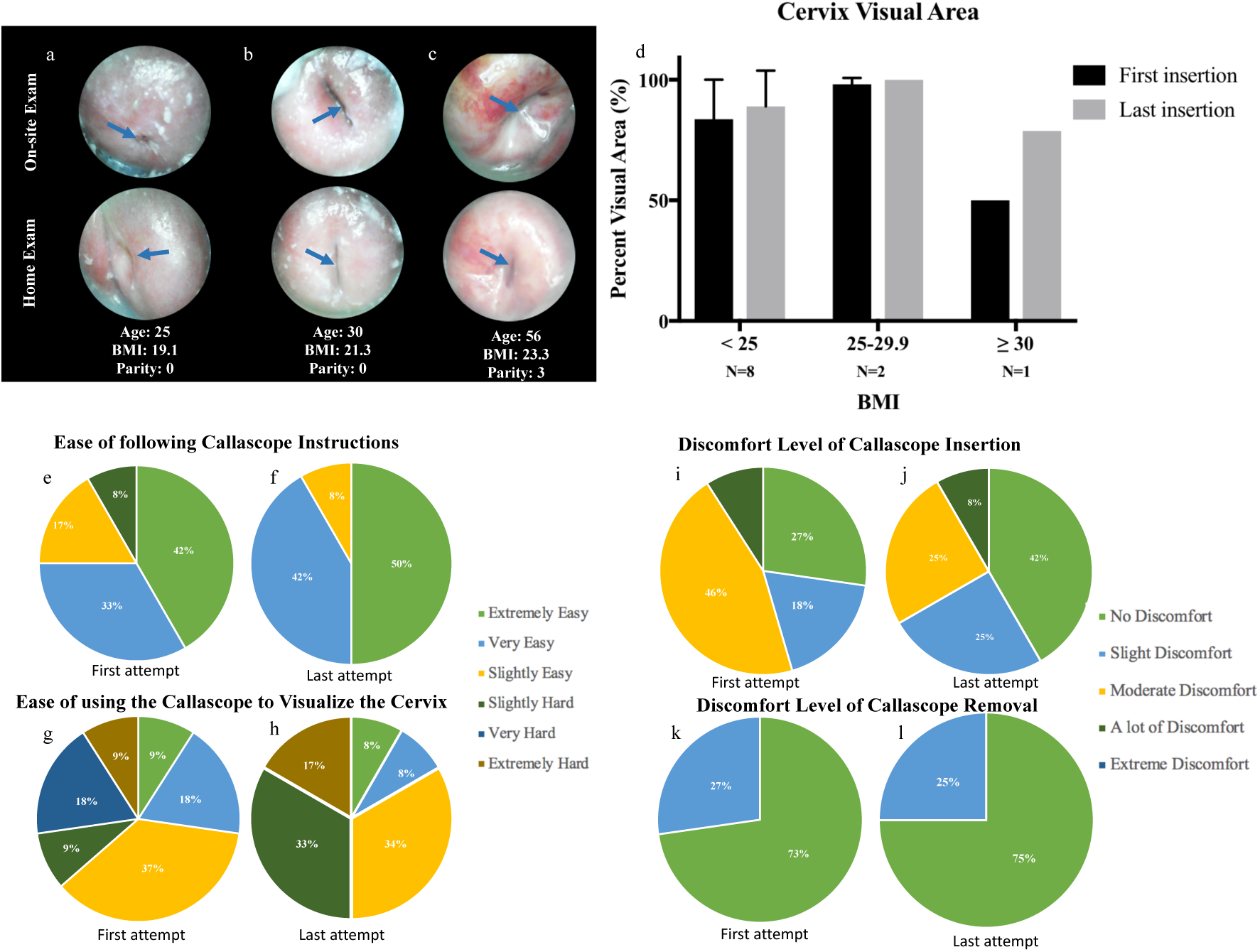
Representative images captured by participants using the Callascope during the on-site exam and the home-exam, respectively. a) Case where participant adequately captured cervix image with centered os in both on-site and home exams, b) case where participant could not center cervical os on-site but was able to do so in the home exam, c) case where participant was able to center os in on-site exam but was unable to do so in-home exam. Blue arrows indicate the cervical os. d) Bar graphs comparing images from first insertion with final insertion. Overall the images don’t show any significant differences in cervix area visualization between the first insertion on-site exam and final insertion at home exam. Error bars are standard deviation, e-l) Responses from post-insertion surveys for first and last attempts of self-exam. e/f) Discomfort level of insertion of the Callascope during first and last use. g/h) Discomfort level of removal of the Callascope during first and last use. i/j) Ease of following the Callascope training instructions during first and last use. k/l) Ease of using the Callascope to find the cervix during first and last use.

### Audio reflections and optional comments provided more in-depth reviews by participants

During the Clinical studies, patients who participated in the study provided optional comments on the experience. Participants from the home study also provided audio reflections of their experience with the Callascope. The themes that emerged from the comments after physician-based exam, and audio reflections after the self-exam, related to comfort, ease of use, visualization, and awareness. (**Table 4**). Almost all of participants mentioned the device was more comfortable than the standard of care speculum. Self-exam participants mentioned that the device was easy to use at home and expressed excitement over being able to view their cervix themselves. Participants also mentioned feeling empowered and having improved cervix awareness by being able to visualize their reproductive anatomy themselves, which they did not have previously. Key words associated with use of the Callascope include “comfortable”, “tampon-like”, “easy to use”, “empowering”, and “fascinating”.

**Table 4:** Themes and representative quotes from clinical study comments and home study audio reflections on experience with Calla exam.

| Emerging themes | % of Participants | Representative Quotes |
| --- | --- | --- |
| Most participants found it easy and comfortable to use the device | 90.9% | “I really liked this device. I thought it was really easy to use. I thought the instructions were really straight-forward. I think it is a really cool idea”<br>“Much better (than the speculum), I like it. I would like this new one every time.”<br>“The inserter was a smoother transition (than the speculum). It felt like someone inserting you with an object literally but with little to no discomfort.”<br>“The new inserter was much more comfortable and easier. It’s never enjoyable going to a Dr., however this is much more welcoming than the old speculum.”<br>“It’s awesome.” |
| About half of the participants mentioned that it was very easy to visualize the cervix at home | 45.45% | “At home I was able to find it [the cervix] very easily, it maybe took me what felt like maybe three minutes at the most “<br>“My Calla exam was very easy. It was very useful to actually be able to see my cervix”<br>“Physicians have a hard time locating my cervix. I did it in 5 min.” |
| Most participants found the experience empowering. | 63.63% | “It was an empowering and intimate experience with myself.”<br>“It’s not something that I have ever thought about before, so it was empowering in that way. |
| Most participants found the experience allowed self-exploration and improved awareness of female reproductive anatomy. | 72.7% | “The pictures were, well, sort of fascinating to see yourself the way a Doctor or a medical student sees other people’s bodies. Some Other feeling you know? Some feeling of alienation in looking at myself in terms of such a physical view that is so different than self-image. Yet also it's just nice to be able to really access your own body visually... It probably left me more motivated to undergo gynecological exams if recommended.”<br>“I’m a cis female in my late twenties with an IUD, so I’ve had several gynecological exams over the years and I have been on the receiving end of a speculum and so I guess I was expecting this to be a similar experience, very clinical and impersonal in a way. I think when I think of female reproductive anatomy I think of that it’s enigmatic, that it’s hidden from us physically. We don’t have a lot of external genitalia, but it’s also kind of hidden in the way that we don’t talk much about vaginas and uteruses and cervixes. So this was kind of an eye-opening experience to be able to visualize something that is so intrinsically a part of me that I have never seen in this way before. It was much more personal and fascinating than I expected it to be.”<br>“I think this is a very cool project and I’m excited to see where it goes and if it does indeed prove useful in both low-resource settings and traditional western higher resource clinics. I think it is a potentially a very useful tool.” |

## Discussion

We have developed the Callascope, a low-cost (∼$50) cervical visualization device that is amenable to physician-based exam or self-exam of the cervix as an alternative to visualization through the duckbill speculum. Both studies demonstrated adequate visualization of the cervix in greater than 80% of participants. Diversity in BMI, vaginal births and age were key to our study since these typically affect speculum-based cervical examinations. Higher BMIs have been associated with vaginal wall prolapse, which causes the walls to obscure the cervix in a speculum-based exam. Number of vaginal births affect the size of the cervix and could affect positioning and visualization. Age-related vaginal atrophy can also influence visualization of the transformation zone. High BMI and vaginal parity were attributed to the inability of the Callascope to visualize the cervix in 20% of the participants and could be potentially addressed by having an extended lip or larger opening designing on the inserter of the Callascope. Patients indicated a significantly greater comfort with the Callascope compared to the speculum. Self-imaging studies demonstrated that women could adequately visualize their cervix with the os centered and without physician guidance using a tutorial guide. While 83% of the participants adequately imaged their cervix on the first try, all the participants eventually imaged their cervix during subsequent tries (usually the second or third). Even though there is the argument that participants in the home study were already self-motivated to seek an alternative to the speculum since they voluntarily signed up to be in the study, participants in the clinical study were patients who were approached the day of their clinic visit and did not have prior motivation to seek out the speculum alternative. Other than our preliminary published work in a pilot group of volunteers for physician-assisted self-imaging of the cervix, there have been no other studies that we know of examining the feasibility for self-imaging of the cervix. The results from this study agree with results from our pilot study in which over 90% of participants mentioned overall preference for the Callascope over the speculum. Our previous results also found that about 83% of users were able to view their cervix with physician assistance, consistent with our new results. ^30^ Future iterations of the device will include a channel for contrast agent administration and mucus removal.

With these studies demonstrating an overwhelming participant preference for the Callascope for self-use and clinical use, the Callascope could provide a patient-centric alternative to the standard speculum in clinical exams and provide an avenue for women to perform basic cervical exams in the comfort of their homes, analogous to the self-breast exam. Studies for self-cervical cancer screening in general have primarily been around the acceptability and feasibility of HPV self-sample collection methods in which women insert a brush into the vagina and rotate it to acquire samples. Acceptability for these studies have been measured by scores of discomforts, pain, embarrassment, and privacy. Even though physician-collected HPV samples were initially viewed as the gold standard, with large scale studies, it has come to light that women are capable of collecting viable samples for HPV screening, and this is currently being recommended for use in several resource-limited settings.^33-35^

Additionally, the unique ability of the Callascope to enable low-cost, comfortable and portable cervical visualization presents an opportunity for its use in educational initiatives to improve women’s awareness of their vaginal and cervical anatomy. While having access to one’s cervix and reflecting on the factors that influence one’s experience of reproductive anatomy in and of itself is insufficient to solve the numerous challenges of female reproductive health, it can be an important and enriching step for many women. For example, in many parts of the world, the lack of cervical cancer awareness is compounded by lack of awareness of the cervix and/or stigma associated with thinking and talking about it. We believe that demystifying the cervix for women and having them engage with their own bodies in a new way can shift the narrative from invisible and negative to visible and positive. We envision that as women discover their own cervices, they will be more comfortable talking about their reproductive health. Increased knowledge of their own cervices from visualization of its anatomy, can improve women’s awareness of and interest in cervical cancer screening and can also improve confidence when self-sampling for cervical cancer screening. The Callascope can also be easily used by transgender men who still have female anatomies yet face various barriers to reproductive health care and education. This could not only serve as a home-based self-screening tool for these men, but also provide awareness of their reproductive anatomy and reduce stigma associated with it.

Overall, the Callascope allows women-centered cervix imaging to reduce barriers to cervical cancer screening, primarily by removing the speculum and associated pain from initial screening, enabling self-screening and providing a tool for women to educate themselves on their reproductive anatomy. The Callascope has the potential to reduce loss to follow-up rates, encourage cervical cancer screening and reduce mortality from the disease. Since it enables cervix visualization, it can also potentially be used by women in the comfort of their homes to view the cervix and vaginal walls for infections, IUD strings and labor dilations. The Callascope is part of a paradigm where technology extends from improving clinic-based medical care to giving women greater control and information at home.

## Conclusion

We have developed the Callascope for comfortable, low-cost and portable visualization of the cervix. The Callascope enables clinical and self-imaging of the cervix and is more accessible and comfortable than the speculum. We believe that the Callascope can also be an important educational tool to help increase awareness of cervical health and cervical cancer screening.

## Acknowledgements

The authors would like to thank Mr. Eric Stach (Department of Mechanical Engineering—Educational Laboratory) and Mr. Christopher Lam for their technical assistance in the fabrication of prototypes and Jennifer Gallagher, Bonnie Thiele, Kristin Weaver, as well as the residents and nurses at the Duke perinatal unit and the Duke obstetrics and gynecology department for their invaluable efforts in support of the home-based and clinical studies.

## Author Contributions

1. **Conceptualization:** MNA JSA MSK RJP MJH GS JSS DJ WH NR.
2. **Data curation:** MNA JSA MED JG BT JWS
3. **Data analysis:** MNA MED
4. **Funding acquisition:** MNA JSA MSK MJH GS DJ WH NR.
5. **Investigation:** MNA JSA MED MSK JG BT JWS NR.
6. **Methodology:** MNA JSA MED MSK JG JWS RJP BT MJH GS JSS DJ WH NR.
7. **Project administration:** MSK JG BT WH NR.
8. **Resources:** MSK WH NR.
9. **Supervision:** MNA MSK JG JWS BT MJH GS JSS DJ WH NR.
10. **Writing – original draft:** MNA MED NR.
11. **Writing – review & editing:** MNA JSA MED MSK JG JWS RJP BT MJH GS JSS DJ WH NR.

## References

1. Comprehensive cervical cancer control: A guide to essential practice. World Health Organization publication 2006. WHO Press WHO, 20 Avenue Appia, 1211 Geneva 27, Switzerland.

2. World Health Organization. Comprehensive cervical cancer prevention and control: a healthier future for girls and women. Geneva: World Health Organization, 2013.

3. WHO Guidelines for Screening and Treatment of Precancerous Lesions for Cervical Cancer Prevention. Geneva; 2013.

4. Hoyo C, Yarnall KS, Skinner CS, Moorman PG, Sellers D, Reid L. Pain predicts non-adherence to pap smear screening among middle-aged African American women. Prev Med 2005; 41(2): 439–45.

5. Khan M, Sultana SS, Jabeen N, Arain U, Khans S. Visual inspection of cervix with acetic acid: a good alternative to pap smear for cervical cancer screening in resource-limited setting. J Pak Med Assoc 2015; 65(2): 192–5.

6. Larsen M, Oldeide CC, Malterud K. Not so bad after all…, Women’s experiences of pelvic examinations. Fam Pract 1997; 14(2): 148–52.

7. Seehusen DA, Johnson DR, Earwood JS, et al. Improving women’s experience during speculum examinations at routine gynaecological visits: randomised clinical trial. BMJ 2006; 333(7560): 171.

8. Kahn JA, Goodman E, Huang B, Slap GB, Emans SJ. Predictors of Papanicolaou smear return in a hospital-based adolescent and young adult clinic. Obstet Gynecol 2003; 101(3): 490–9.

9. Wright D, Fenwick J, Stephenson P, Monterosso L. Speculum ‘self-insertion’: a pilot study. J Clin Nurs 2005; 14(9): 1098–111.

10. Lyimo FS, Beran TN. Demographic, knowledge, attitudinal, and accessibility factors associated with uptake of cervical cancer screening among women in a rural district of Tanzania: three public policy implications. BMC Public Health 2012; 12: 22.

11. Crowley D LG. Emotional aspects of gynecology: depression, anxiety, PTSD, eating disorders, substance abuse, “difficult” patients, sexual function, rape, intimate partner violence, and grief. Comprehensive Gynecology. 6 ed; 2012.

12. Garcia-Moreno C, Jansen HA, Ellsberg M, et al. Prevalence of intimate partner violence: findings from the WHO multi-country study on women’s health and domestic violence. Lancet 2006; 368(9543): 1260–9.

13. Williams CM, McCloskey LA, Larsen U. Sexual violence at first intercourse against women in Moshi, northern Tanzania: prevalence, risk factors, and consequences. Popul Stud (Camb) 2008; 62(3): 335–48.

14. (IARC) IAfRoC. GLOBOCAN Cervix Uteri ASR (W) per 100,000, all ages. 2012. http://globocan.iarc.fr/old/bar_sex_site.asp?selection=4162&title=Cervix+uteri&statistic=2&populations=6&window=1&grid=1&color1=5&color1e=&color2=4&color2e=&submit=%C2%A0Execute (accessed 27 June 2016).

15. Ostensson E, Alder S, Elfstrom KM, et al. Barriers to and facilitators of compliance with clinic-based cervical cancer screening: population-based cohort study of women aged 23-60 years. PLoS One 2015; 10(5): e0128270.

16. Yoo W, Kim S, Huh WK, et al. Recent trends in racial and regional disparities in cervical cancer incidence and mortality in United States. PLoS One 2017; 12(2): e0172548.

17. Franco EL. Self-sampling for cervical cancer screening: Empowering women to lead a paradigm change in cancer control. Curr Oncol 2018; 25(1): e1–e3.

18. Musa J, Achenbach CJ, O’Dwyer LC, et al. Effect of cervical cancer education and provider recommendation for screening on screening rates: A systematic review and meta-analysis. PLoS One 2017; 12(9): e0183924.

19. Jacobson M. Everything You Need to Know About Vaginas. Women’s Health. 2014.

20. Fylan F. Screening for cervical cancer: a review of women’s attitudes, knowledge, and behaviour. Br J Gen Pract 1998; 48(433): 1509–14.

21. Shedlin M, Amastae J, Potter JE, Hopkins K, Grossman D. Knowledge and beliefs about reproductive anatomy and physiology among Mexican-Origin women in the USA: implications for effective oral contraceptive use. Cult Health Sex 2013; 15(4): 466–79.

22. El Gelany S, Moussa O. Reproductive health awareness among educated young women in Egypt. Int J Gynaecol Obstet 2013; 120(1): 23–6.

23. Jayant K, Rao RS, Nene BM, Dale PS. Improved stage at diagnosis of cervical cancer with increased cancer awareness in a rural Indian population. Int J Cancer 1995; 63(2): 161–3.

24. Suarez L, Roche RA, Nichols D, Simpson DM. Knowledge, behavior, and fears concerning breast and cervical cancer among older low-income Mexican-American women. Am J Prev Med 1997; 13(2): 137–42.

25. Sankaranarayanan R, Budukh AM, Rajkumar R. Effective screening programmes for cervical cancer in low- and middle-income developing countries. Bull World Health Organ 2001; 79(10): 954–62.

26. Patra S, Upadhyay M, Chhabra P. Awareness of cervical cancer and willingness to participate in screening program: Public health policy implications. J Cancer Res Ther 2017; 13(2): 318–23.

27. Eze JN, Umeora OU, Obuna JA, Egwuatu VE, Ejikeme BN. Cervical cancer awareness and cervical screening uptake at the Mater Misericordiae Hospital, Afikpo, Southeast Nigeria. Ann Afr Med 2012; 11(4): 238–43.

28. Coronado Interis E, Anakwenze CP, Aung M, Jolly PE. Increasing Cervical Cancer Awareness and Screening in Jamaica: Effectiveness of a Theory-Based Educational Intervention. Int J Environ Res Public Health 2015; 13(1): ijerph13010053.

29. Arevian M, Noureddine S, Kabakian-Khasholian T. Raising awareness and providing free screening improves cervical cancer screening among economically disadvantaged Lebanese/Armenian women. J Transcult Nurs 2006; 17(4): 357–64.

30. Asiedu MN, Agudogo J, Krieger MS, et al. Design and preliminary analysis of a vaginal inserter for speculum-free cervical cancer screening. PLoS One 2017; 12(5): e0177782.

31. Administration FaD. Cleared Sterilants and High Level Disinfectants with General Claims for Processing Reusable Medical and Dental Devices. 2015. http://www.fda.gov/MedicalDevices/DeviceRegulationandGuidance/ReprocessingofReusableMedicalDevices/ucm437347.ht m.

32. William A. Rutala PD M.P.H., David J. Weber, M.D. M.P.H. and the Healthcare Infection Control Practices Advisory Committee (HICPAC). Guideline for Disinfection and Sterilization in Healthcare Facilities. Center for disease control 2008.

33. Waller J, McCaffery K, Forrest S, et al. Acceptability of unsupervised HPV self-sampling using written instructions. J Med Screen 2006; 13(4): 208–13.

34. Petignat P, Faltin DL, Bruchim I, Tramer MR, Franco EL, Coutlee F. Are self-collected samples comparable to physician-collected cervical specimens for human papillomavirus DNA testing? A systematic review and meta-analysis. Gynecol Oncol 2007; 105(2): 530–5.

35. Dzuba IG, Diaz EY, Allen B, et al. The acceptability of self-collected samples for HPV testing vs. the pap test as alternatives in cervical cancer screening. J Womens Health Gend Based Med 2002; 11(3): 265–75.

